# Active information maintenance in working memory by a sensory cortex

**DOI:** 10.1101/385393

**Authors:** Xiaoxing Zhang, Wenjun Yan, Wenliang Wang, Hongmei Fan, Ruiqing Hou, Yulei Chen, Zhaoqin Chen, Shumin Duan, Albert Compte, Chengyu T. Li

**Affiliations:** institute of Neuroscience, State Key Laboratory of Neuroscience, Key Laboratory of Primate Neurobiology, CAS Center for Excellence in Brain Science and Intelligence Technology, Shanghai Institutes for Biological Sciences, Chinese Academy of Sciences, Shanghai 200031, China; School of Future Technology, University of Chinese Academy of Sciences, Beijing 100049, China; Department of Neurobiology, Key Laboratory of Medical Neurobiology of the Ministry of Health of China, Key Laboratory of Neurobiology, Zhejiang University School of Medicine, Hangzhou, Zhejiang, 310058, China; Institut d’Investigacions Biomédiques August Pi i Sunyer, 08036 Barcelona, Spain

**Keywords:** Working memory, dual-task, delay-period activity, anterior piriform cortex, optogenetics, population neural activity

## Abstract

Working memory is a critical function of the brain to maintain and manipulate information over delay periods of seconds. Sensory areas have been implicated in working memory; however, it is debated whether the delay-period activity of sensory regions is actively maintaining information or passively reflecting top-down inputs. We hereby examined the anterior piriform cortex, an olfactory cortex, in head-fixed mice performing a series of olfactory working memory tasks. Information maintenance is necessary in these tasks, especially in a dual-task paradigm in which mice are required to perform another distracting task while actively maintaining information during the delay period. Optogenetic suppression of the piriform cortex activity during the delay period impaired performance in all the tasks.Furthermore, electrophysiological recordings revealed that the delay-period activity of the anterior piriform cortex encoded odor information with or without the distracting task.Thus, this sensory cortex is critical for active information maintenance in working memory.

## Introduction

Working memory (WM) is a critical function of the brain to actively maintain and manipulate information over a delay period of several seconds (Baddeley, 2012).A buffer between recent external inputs and immediate behavioral outputs, WM is a critical component of cognition (Jonides et al., 2008). Originally the prefrontal cortex was thought to be the site for WM (Jacobsen, 1935).However, later results suggested that parallel distributed circuits are responsible for WM (Christophel et al., 2017; D’Esposito and Postle, 2015; Eriksson et al., 2015). Previously we have shown that the delay-period activity of the medial prefrontal cortex (mPFC) of mice is only important during the learning but not the well-trained phase in an olfactory WM task (Liu et al., 2014).Here, we seek to elucidate the contribution of a sensory cortex to WM after mice are well-trained.

Sensory regions exhibit the delay-period activity that can code maintained information (Fuster and Jervey, 1981; Harrison and Tong, 2009; Mendoza-Halliday et al., 2014; Miyashita and Chang, 1988; Pasternak and Greenlee, 2005).Furthermore, perturbation of neural activity in sensory areas can impair WM performance (Colombo et al., 1990; Guo et al., 2014; Harris et al., 2002; Pasternak and Greenlee, 2005; Seidemann et al., 1998).However, these previous perturbation experiments either lacked the temporal specificity in the perturbation methods (Fenno et al., 2011), or mixed WM maintenance with sensory perception or decision choices in the behavioral design (Passingham and Wise, 2012).Additionally, single-neuron recording studies sometimes failed to find robust memory signals in early sensory cortices (Hernandez et al., 2010).Moreover, it has been debated whether the delay-period activity of sensory regions is actively maintaining information (Pasternak and Greenlee, 2005) or passively reflecting higher associative areas via top-down inputs (Mendoza-Halliday et al., 2014; van Kerkoerle et al., 2017). These two theories make different predictions. The active, but not the passive, theory predicts behavioral deficits when perturbing the delay-period activity in sensory regions, as well as sustained and informative delay-period activity in sensory areas robust to distractors.Indeed, a key argument for the passive top-down theory is that the delay-period activity of the sensory cortices degrades in the presence of distractors, whereas the activity of the fronto-parietal cortices remains robust (Bettencourt and Xu, 2016; Miller et al., 1996).

In the current study we tackled this debate in the anterior piriform cortex (APC) for olfactory WM. The APC is a good candidate because it is directly connected with the olfactory bulb (Bekkers and Suzuki, 2013; Davison and Ehlers, 2011; Ghosh et al., 2011; Miyamichi et al., 2011; Price and Powell, 1970; Sosulski et al., 2011), encodes odor information (Bekkers and Suzuki, 2013; Courtiol and Wilson, 2016; Gire et al., 2013; Iurilli and Datta, 2017; Miura et al., 2012; Poo and Isaacson, 2009; Schoenbaum and Eichenbaum, 1995; Shusterman et al., 2011; Stettler and Axel, 2009; Wilson, 1998; Zhan and Luo, 2010), and is important for olfactory behavior (Bekkers and Suzuki, 2013; Choi et al., 2011; Courtiol and Wilson, 2016; Miura et al., 2012). The rich recurrent connectivity within the APC (Bekkers and Suzuki, 2013; Franks et al., 2011) is also well suited for generating reverberating activity that outlasts sensory inputs (Major and Tank, 2004). In addition, activity patterns of the APC neurons are critically dependent on olfactory experience (Courtiol and Wilson, 2016; Roesch et al., 2007; Saar et al., 1999; Wilson, 1998).Finally, the piriform cortex is also activated in a human olfactory WM task (Zelano et al., 2009). We trained head-fixed mice to perform a series of olfactory WM tasks, with or without a distracting task during the delay period. We found that optogenetic suppression of the APC activity during the delay period impaired performance in all the tasks.Furthermore, electrophysiological recordings revealed that the delay-period activity can code odor information with or without the distracting task.Thus, the APC is critical for active information maintenance in olfactory working memory.

## Results

### The design and performance of a delayed non-match to sample task

To temporally dissociate information maintenance from perception and decision making, we firstly trained head-fixed mice to perform an olfactory delayed non-match to sample (DNMS) task (Liu et al., 2014), using an automatic training system (Han et al., 2018). In this task, two odors (S1, n-butanol; or S2, propyl formate) were presented to mice, separated by a delay period (5-40 sec, **Figures 1A and 1B, Movie S1**). Licking following the non-match trials (S1-S2, or S2-S1) lead to water reward. No explicit punishment was applied for error trials. Mice readily performed the task with little licking during the delay period (**Figure 1C**).A hallmark of WM is the progressive decay of performance with increasing delay duration (Baddeley, 2012), which was indeed observed in the task (**Figure 1D**).

**Figure 1.**
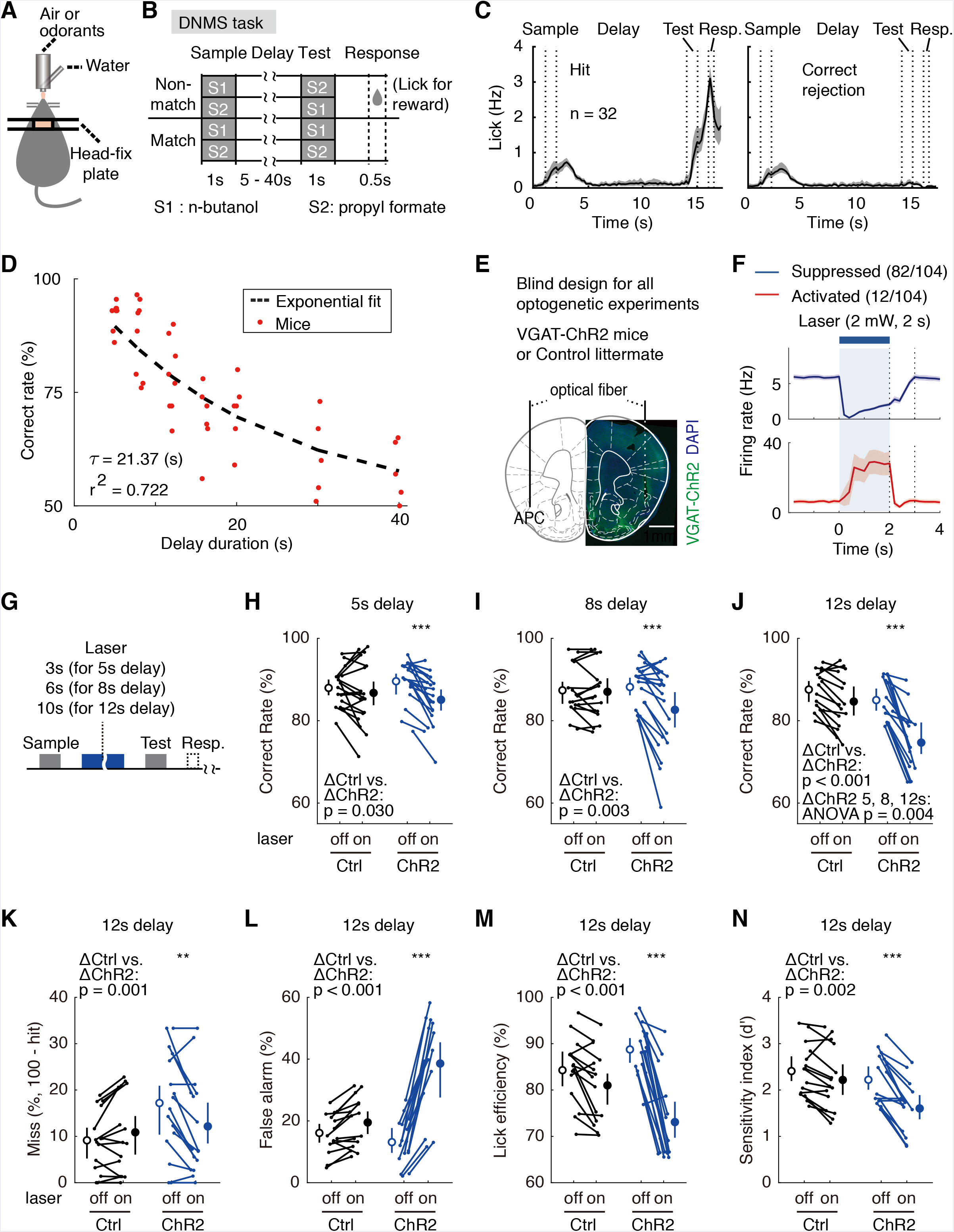
The APC activity is important for the DNMS performance. (**A**) Diagram of the experimental setup.(**B**) Design of the DNMS task.(**C**) Averaged licking rate of mice well-trained for the DNMS task in hit (left) and correct rejection (right) trials.(**D**) The DNMS task correct rate with varied delay durations, fitted to an exponential-decay function.(**E**) Targeted region and sites of optical fiber insertion.(**F**) Effectiveness of optogenetic silencing by awake op-tetrode recording in *vivo*.(**H-J**) Correct rates in the DNMS task following the APC delay-period optogenetic suppression. Statistic inset, the Wilcoxon Rank-Sum test; numbers above ChR2 groups, ***, p < 0.001, Student’s paired t-test; error bars: 95% confidence interval of the mean, unless stated otherwise. Delay duration, 5s (**H**), 8s (**I**) and 12s (**J**).(**K-N**) Miss rate, false alarm rate, lick efficiency and sensitivity index (d’) following the APC delay-period suppression in the DNMS task with 12s delay.

### The APC delay-period activity is critical in olfactory DNMS

It has previously been shown that the delay-period activity of the medial prefrontal cortex (mPFC) is important only during learning of the DNMS task, but not after mice were well-trained (Liu et al., 2014). To test the potential role of the APC in the well-trained phase, we optogenetically (Fenno et al., 2011) suppressed the delay-period activity of the APC pyramidal neurons after mice were well-trained (correct rate above 80% in 40 consecutive trials). Transgenic mice expressing channel-rhodopsin (ChR2) only in the GABAergic neurons [with the promoter of vesicular GABA transporter (VGAT), referred to as VGAT-ChR2, (Zhao et al., 2011)] were used as the experimental group, whereas the ChR2-negative littermates were used in the control group in a blind design (**Figure 1E**, see STAR Methods). The expression and effectiveness of ChR2 were verified by immunostaining and op-tetrode recording, respectively (**Figure 1F**). Activity suppression was achieved by step-laser illumination (473 nm, 2 mW) during the laser-on trials, each followed by a laser-off trial (**Figure 1G**). Performance was impaired following optogenetic suppression of the APC delay-period activity in the VGAT-ChR2 mice, with delay duration of 5, 8, or 12 sec (**Figures 1H-1J**).Moreover, optogenetic suppression in the trials with longer delay duration resulted in larger deficits (**Figures 1H-1J**). The behavioral defects were further manifested as the increased false alarm rate and decreased discriminability *d’* (false alarm rate for all delay duration, *d’* for 8s and 12s delay duration, **Figures 1L, 1N, S1B, S1F and S1H**).Similarly, the lick efficiency (the percentage of the successful licks with rewards) was impaired in the laser-on trials in the VGAT-ChR2 mice (**Figures 1M, S1C and S1G**).

The optogenetic manipulation during the delay period might impair sensory perception during the second-odor delivery period. To exclude this possibility, we suppressed the APC activity before odor delivery in the following three tasks. In all three tasks, we observed the intact behavioral performance (**Figure 2A**).Firstly, we optogenetically suppressed the APC activity before the sample-delivery period (baseline) in the DNMS task (**Figure 2B**).Secondly, mice were trained to perform a sensory-discriminating Go/No-go (GNG) task. The APC activity before the test odor was optogenetically suppressed (**Figure 2C**).Thirdly, mice were trained to perform a non-match-to-sample without delay (NMS-WOD) task. The APC activity before and after test-odor delivery was suppressed in the tasks (**Figure 2D**). The lack of behavioral impairment in all three control experiments excluded the perception-impairment hypothesis. Therefore the functional role of the APC delay activity should be information maintenance.

**Figure 2.**
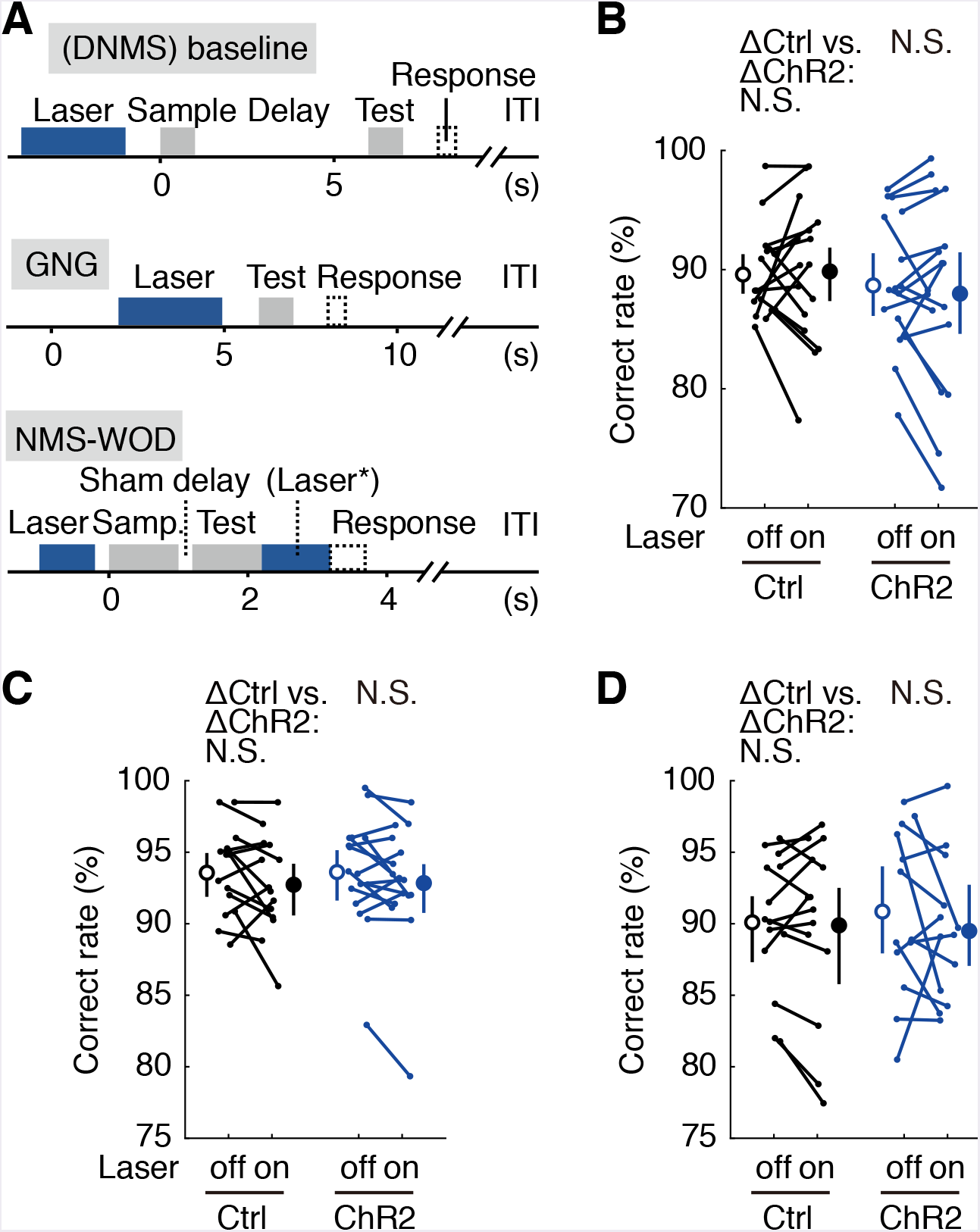
The optogenetic perturbation effects were not due to impaired sensory perception. (**A**) Designs of the DNMS-baseline perturbation, Go-Nogo (GNG) and non-match to sample without delay (NMS-WOD) optogenetic control tasks.(**B-D**) Correct rates in all three control tasks. Statistics above control groups, the Wilcoxon Rank-Sum test; statistics above ChR2 groups, paired t-test; error bars: 95% confidence interval of the mean, N.S., not significant.

### Formal model comparison further demonstrating the importance of APC delay-period activity

The behavioral defects following optogenetic perturbation of the APC delay-period activity reflected its critical contribution to information maintenance. In order to quantify this contribution, we designed seven candidate generalized linear models (GLMs) by systematically varying the combinations of task parameters in fitting the performance (**Figure 3A**).A good model should exhibit a high R2 in explaining performance and a low Akaike information criterion (AIC, see STAR Methods), which punishes higher number of free parameters. Such formal model comparison can quantitatively reveal relative contribution of different behavioral parameters in determining performance (Brunton et al., 2013; Hwang et al., 2017).

**Figure 3.**
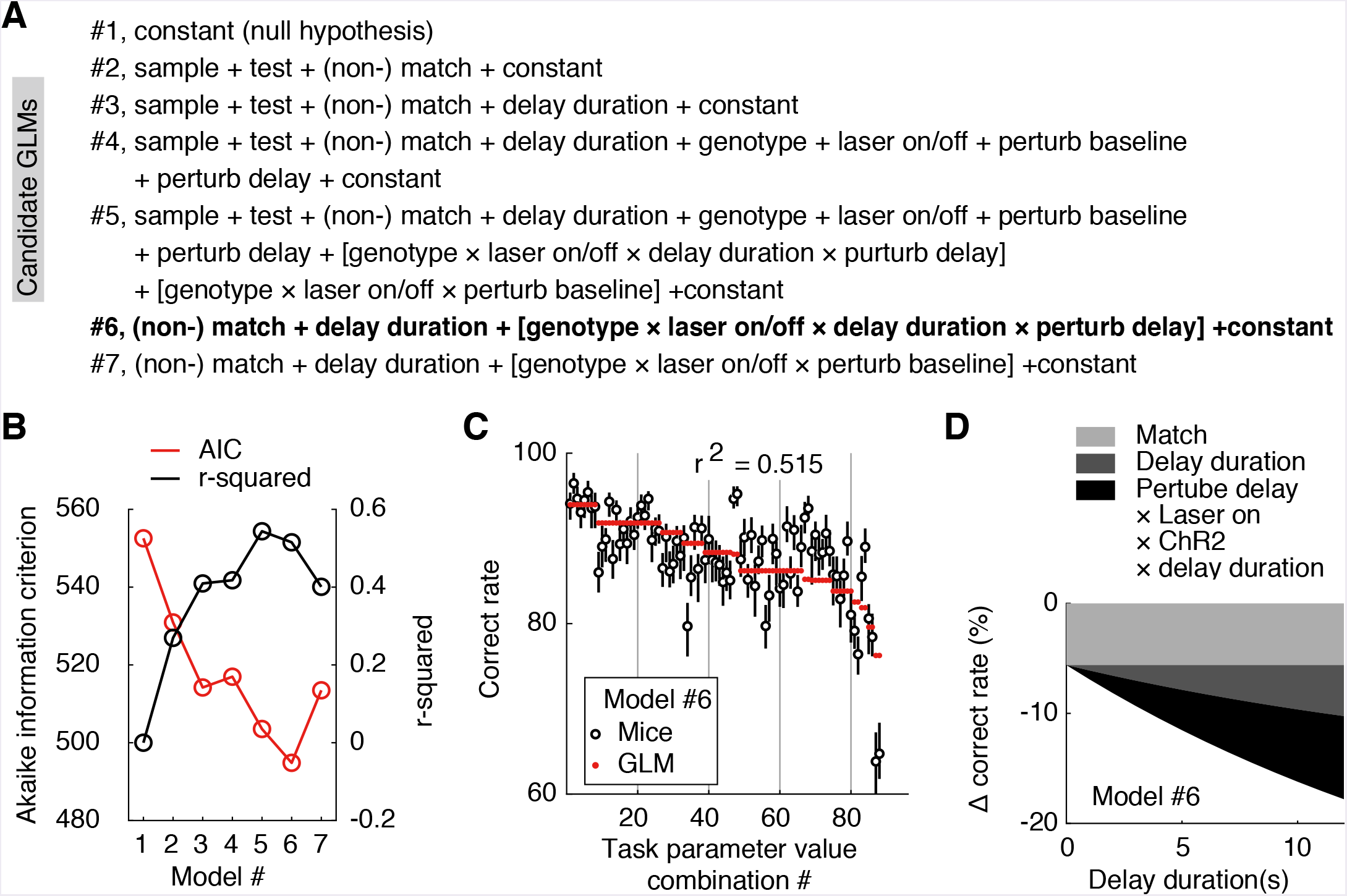
Formal model comparison demonstrating the importance of APC delay-period activity. (**A**) Design of the candidate GLMs. The task parameters in each model were then used to fit task performance. Square brackets, interactions.(**B**) Comparison of the candidate GLMs in terms of AIC and r2 in explaining performance.(**C**) Correlations of the performance of mice in the WM tasks and the optimized model (#6 in A). Error bar, 95% confidence interval of mean from bootstrap.(**D**) Comparison in weights of model parameters in #6, showing the predicted effect size against delay duration.

We started with the null-hypothesis that no task parameter affects performance (#1, **Figures 3A** and **3B**). Adding the variables of both the sensory cues (#2) and the time constant for memory decay (#3, from **Figure 1D**) improved the performance of the models, consistent with the obvious importance of sensory cues and delay duration in this task. The genotypes, laser on/off, and the perturbation in delay/baseline period did not further improve the model if added individually (#4).However, the interaction among all these terms improved the model (#5), consistent with the impaired performance in the optogenetic suppression during the delay-period for the VGAT-ChR2 mice, but not during the baseline period or for the control mice. By eliminating less predictive variables, we obtained the optimized model (#6, **Figures 3A-3C**, **Tables S1** and **S2**), which contained only three key parameters and the interactions among them: optogenetic suppression or not during the delay period, match or non-match relationship, and the delay duration. This association between the WM delay perturbation and the delay durations is consistent with the critical role of the APC delay activity in information maintenance (**Figure 3D**).Quantitatively, the impact of optogenetic perturbation out-weighted that of the delay duration (**Figure 3D**). The above analysis further demonstrated the importance of the APC delay-period activity for WM in the olfactory DNMS task.

### The importance of the APC delay-period activity in an olfactory delayed paired association task

The activity of a sensory cortex can be suppressed or enhanced following the repetition of the sensory stimuli (Miller et al., 1993). It has also been suggested that the DNMS task in monkeys can be performed through a recency effect and with little selective memory maintenance (Wittig and Richmond, 2014). To eliminate the involvement of the repeated sample cues, we trained mice to perform an olfactory delayed paired association (DPA) task (**Figure 4A**). In this task, mice were trained to establish an association between the specific pairs of the sample-test odors separated by a delay period (**Figure 4A**, modified from ref.(Schoenbaum and Eichenbaum, 1995)). In each trial, a sample odor (S1, ethyl acetate; or S2, 3-methyl-2-buten-1-ol) was presented, followed by a delay period (13 sec), then a test odor (T1, n-butanol; or T2, propyl formate). Mice were rewarded with water if they licked within the response window in the paired trials (S1-T1 or S2-T2), but not in the unpaired trials (S1-T2 or S2-T1, **Figure 4A**). Successful performance of the task required WM maintenance and learning of arbitrary association between the odor pairs.Importantly, using repetition (Miller et al., 1993) or recency effect (Wittig and Richmond, 2014) cannot solve the task, because the sample odor is not repeated within a trial. Mice readily learnt the task (**Figure 4D**, the black curve) and optogenetic suppression of the APC delay-period activity impaired the DPA performance (**Figures 4B, S2A-S2D**).Therefore, the APC delay-period activity is critical for WM, even in a task without repetition of the sensory cues.

**Figure 4.**
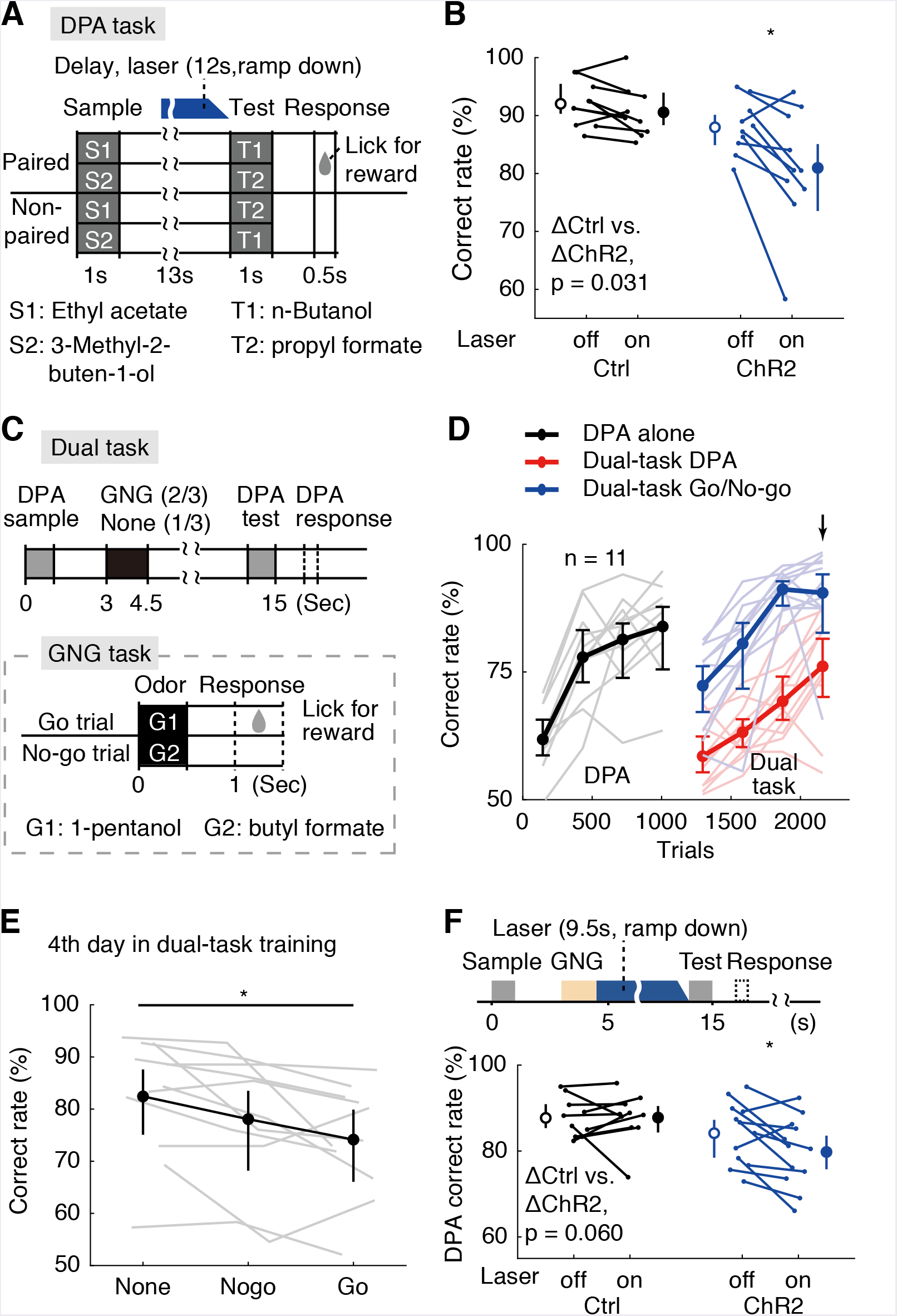
Active memory maintenance by the APC delay activity. (**A**) Design of the delayed paired association (DPA) task.(**B**) Correct rates following the APC delay-period optogenetic suppression in the DPA task. Statistics inset, the Wilcoxon Rank-Sum test; numbers above ChR2 group, Student’s paired t-test, *, p = 0.015.(**C**) Behavioral diagram for the dual-task design.(**D**) Learning curve for the correct rate in the dual-task. Noted the drop of DPA performance after inserting GNG task in the delay period (in red).Arrow, the 4th day in dual-task training.(**E**) Dual-task interference, from performance of 4th day in dual-task training.*, p = 0.041, one-way repeated measure ANOVA.(**F**) Correct rate after suppressing the APC activity during the later-phase delay period after distractors. Statistics inset, the Wilcoxon Rank-Sum test; numbers above ChR2 group, Student’s paired t-test, *, p = 0.021.

### The APC delay activity is critical for active maintenance against a distracting task

A hallmark of active maintenance in WM is resistance against distractors during the delay period (Baddeley, 2012; Bettencourt and Xu, 2016; Miller et al., 1996). To test this ability in mice, we added a distracting GNG task during the delay period of the DPA task (**Figure 4C**). This paradigm belongs to the dual-task designs in studying the central executive control (Baddeley, 2012; Watanabe and Funahashi, 2014), because mice were required to split attention in the middle of the delay period and perform the GNG task, while simultaneously maintaining the sample information of the DPA task (**Figure 4C**). Mice can indeed perform the dual task (**Figure 4D, the red curve**). The DPA performance in the dual-task paradigm was reduced compared to that of the simple DPA task (**Figures 4D** and **4E**), consistent with the dual-task interference observed in human (Baddeley, 2012) and monkeys (Watanabe and Funahashi, 2014).Moreover, interference was dependent on the trial types inserted in the DPA delay, with the worst performance for the Go-distractor trials (**Figure 4E**). We then optogenetically suppressed the delay-period activity of the APC pyramidal neurons after the distracting GNG task. Behavioral performance of DPA task was significantly impaired in the laser-on trials in the VGAT-ChR2 group (**Figure 4F**). The performance in the laser-on vs.laser-off trials between the VGAT-ChR2 and control groups tended to differ (P = 0.06, Rank-sum test). The difference in the false alarm rate in the laser-on and –off trials was statistically significant between the VGAT-ChR2 and control mice (**Figure S2F**). As a negative control, optogenetic suppression before sample delivery did not affect performance (**Figures S2I-S2N**). Therefore the APC delay-period activity is critical for active maintenance against a distracting task.

### Neural correlates of the APC activity in the DNMS task

To investigate the neural correlates of the APC in WM, we recorded the single-unit activity (**Figures 5A** and **5B**) using custom-made tetrodes (**Figure S3**) (Liu et al., 2014). Mice were trained with the DNMS task with a 4-sec delay period and recording started from the first day of the training. Recording electrodes were advanced daily (approximately 50 μm/day after recording sessions). After 2-4 days into the training, the delay duration was increased to 8 sec. We obtained 204 neurons from 18 mice while they performed the task with 4-sec delay duration and 156 neurons from 17 mice with 8-sec delay duration. The activity of many APC neurons in the delay period was related to the identity of the sample odors (an example neuron in **Figure 5A**). Population coding for the maintained information can be visualized in a neuronal-activity heat map, grouped according to the different sample odors (4 and 8 sec in **Figures S3** and **5B**, respectively).A population decoding analysis based on support vector machine (SVM) also demonstrated clear decoding power of the APC delay activity for the sample odors (**Figures 5C** and **5D**).Interestingly, the dynamics of the decoding accuracy in the 8s-delay-tasks closely followed that of the 4s-delay-tasks that scaled-up with a factor of two, consistent with the active maintenance of information in accordance with task requirement.

**Figure 5.**
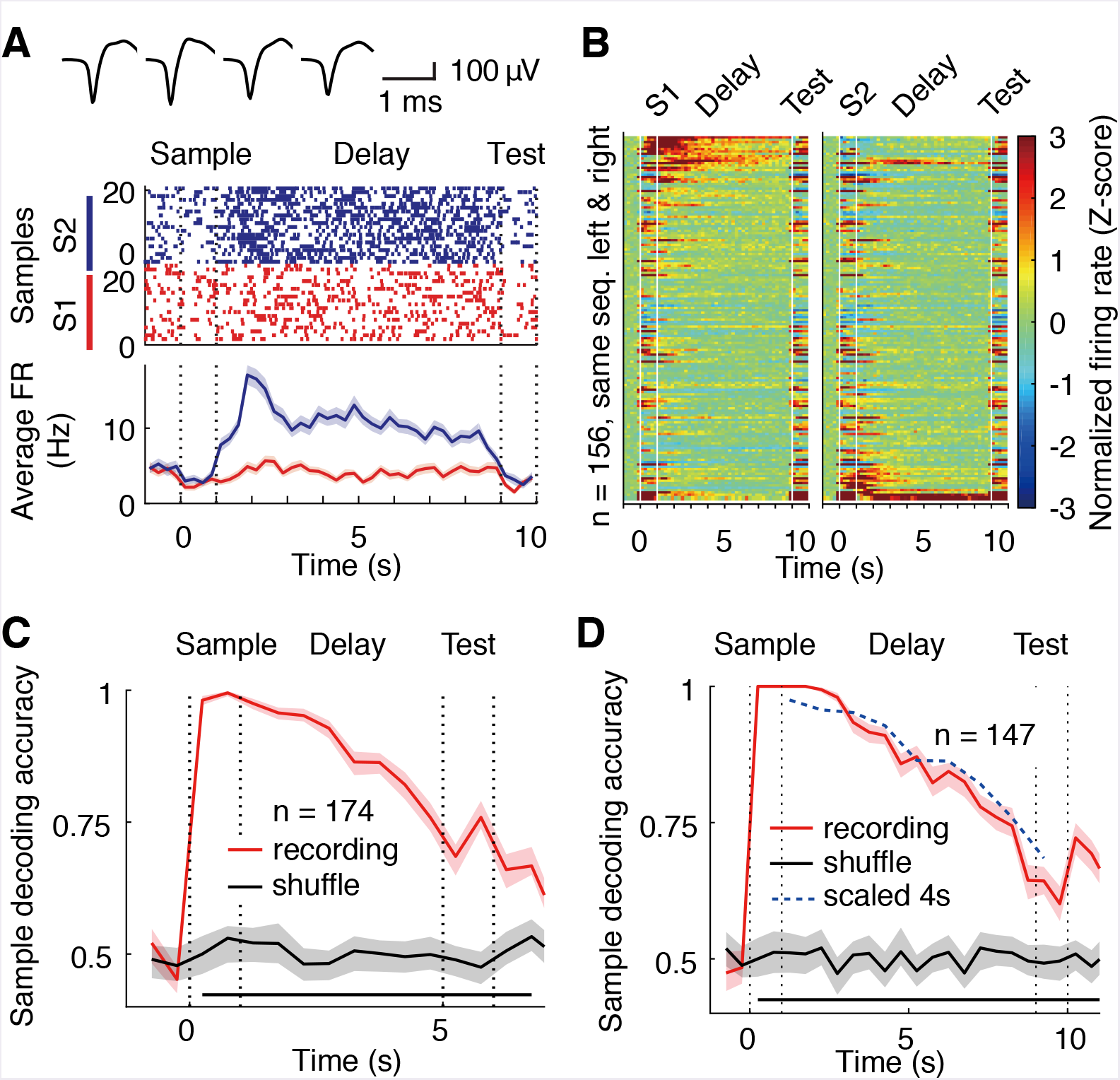
Neural correlates of the APC activity in the DNMS task. (**A**) Spike-raster (above) and peristimulus time histogram (PSTH, below) of an example neuron recorded during the DNMS task. Shadow for PSTH, standard error of the mean (SEM).Top: spike waveforms from tetrode recording.(**B**) Activity of neurons in the DNMS task with odor S1 (left) and S2 (right) as sample.Color: firing rates (FR) in Z-scores. Neurons were aligned according to the FRS1 – FRS2 values during the delay period. White line indicated for the onset of offset of odor delivery.(C and D) Sample decoding accuracy of the APC neuronal activity based on the SVM, in the DNMS task with 4s (**C**) or 8s (**D**) delay.Shadow, 95% confidence interval of mean. Bottom black bar, p < 0.001, permutation test of 1000 repeats, unless stated otherwise.

### Neural correlates of the APC activity in the multi-sample DPA task

It was shown that the number of stimuli used for training can bias the strategies of the animals (Slotnick, 2001), so one might argue about the generality of the results using just two odorants as the sample odors. We therefore trained the mice with a DPA task using six odorants samples (S1-S6) and two odorants as test (T1 and T2), in which S1, S2 and S3 paired with T1, and S4, S5 and S6 paired with T2 for reward (multiple-sample DPA, **Figure 6A**). Despite an initial relearning in each day, mice performed this task well (**Figure 6B**). Similar to the DNMS task with two sample odors, we observed many neurons with sample-odor information during the delay period, as revealed by mutual information (MI, **Figure 6C**, see STAR Methods), which measures to what degree the responses are informative about the identity of the stimulus.A similar results were also obtained with the SVM decoding analysis (**Figure 6D**).

**Figure 6.**
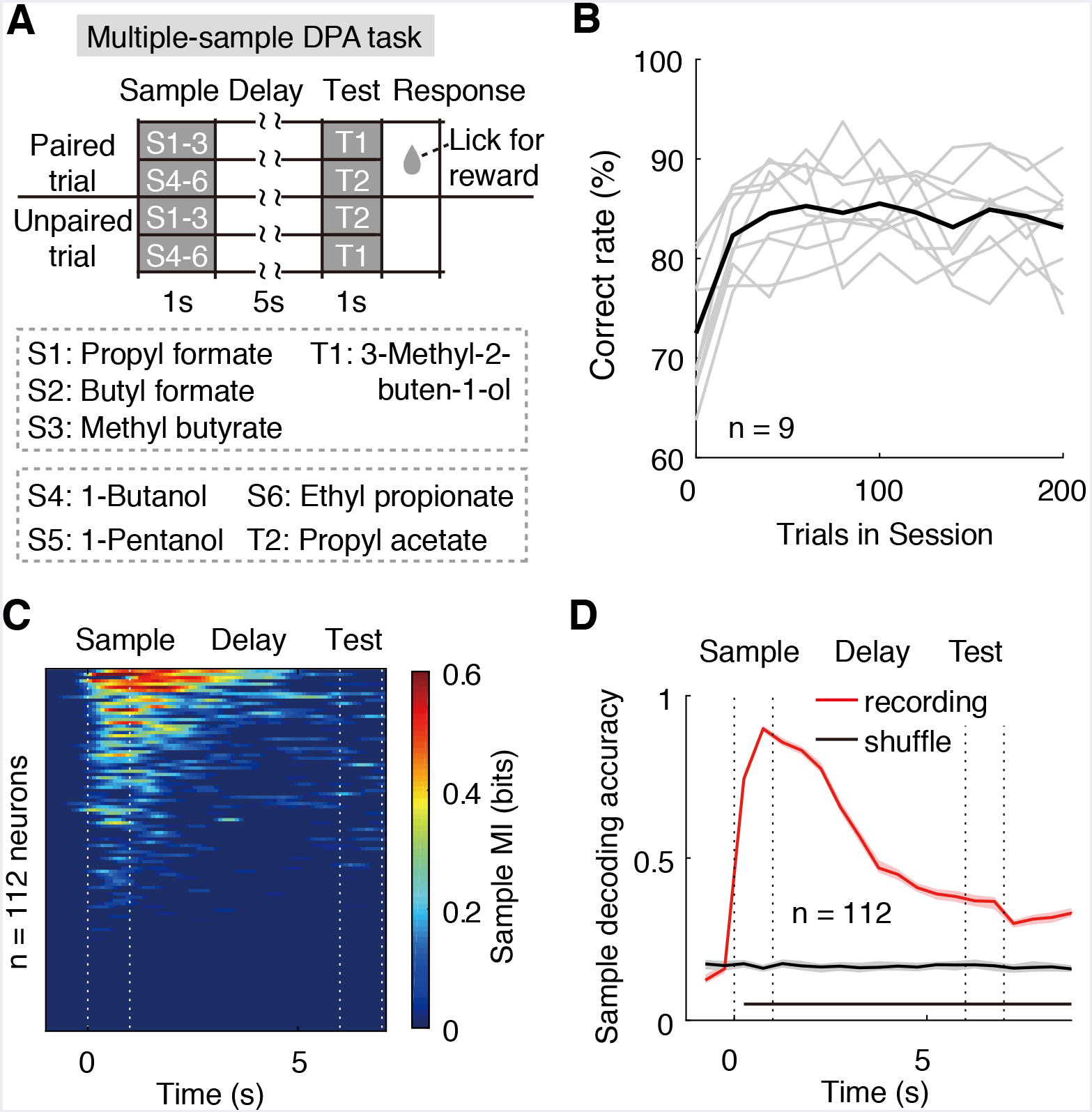
Neural correlates of the APC activity in the multiple-sample DPA task. (**A**) Design of the multiple-sample DPA (MS-DPA) task.(**B**) Averaged daily performance of well-trained mice in the MS-DPA task.(**C**) Mutual information of the APC neurons for the sample odors in the MS-DPA task.(**D**) Sample decoding accuracy of the APC neurons in the MS-DPA task.Shadow, 95% confidence interval of mean. Bottom black bar, p < 0.001, permutation test of 1000 repeats, unless stated otherwise.

### Neural correlates of the APC activity in the dual-task design

We then examined APC neuronal activity in the dual task. Even after application of the distracting task during the delay period, APC population activity still coded the DPA samples and GNG distractors, as shown in the MI analysis of one example neuron in **Figure 7A**. In the APC neuronal population, approximately 30% of neurons can code for both the DPA samples and GNG odors (**Figure 7B**). In the SVM decoding analysis, the APC activity can code for the DPA sample odors in the dual task design without distractor, with the No-go distractor, or with the Go distractor (**Figures 7C-7E**) after the distracting task. As mentioned in the previous optogenetic results, the DPA performance was related to the distractor type in the recording sessions (**Figures 4E and 7F**).Interestingly, we observed that the SVM decoding accuracy during the delay period after the distractor task was also related to the distractor type and performance (**Figure 7F**), supporting the critical role of APC selectivity in performing the DPA task in the dual-task design.

**Figure 7.**
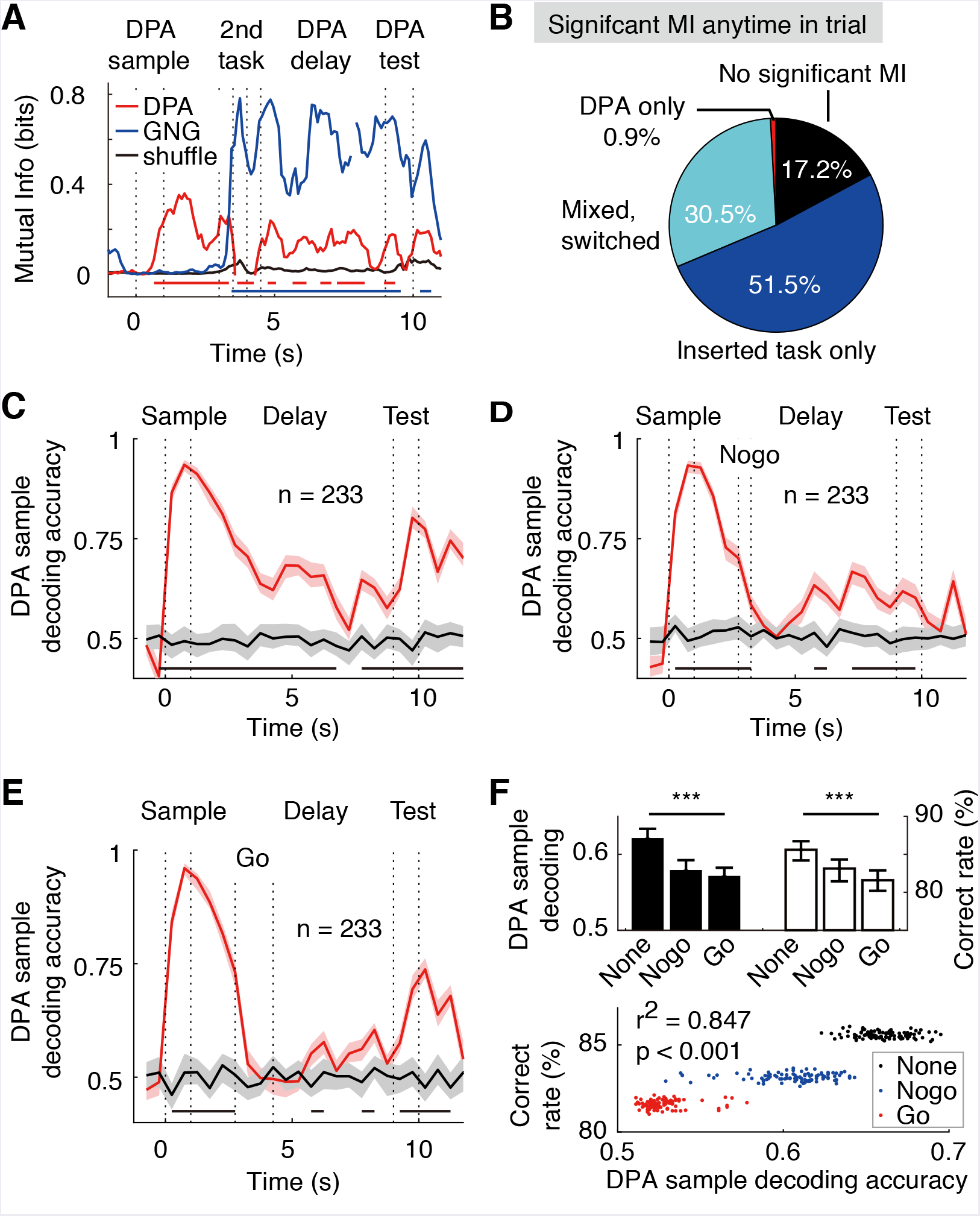
Neural correlates of the APC activity in dual task. (**A**) An example neuron with mixed DPA-sample and GNG-task odor-cue information in the dual-task. Bottom bars, p < 0.001 for DPA sample (red) and GNG cue (blue), permutation test of 1000 repeats, comparing to shuffled control.(**B**) Fraction of neurons with exclusive or mixed/switched odor-cue information in the dual-task.(**C**) DPA sample decoding accuracy of the APC neurons in the dual-task trials without distractor. Bottom black bar, p < 0.001, permutation test of 1000 repeats, unless stated otherwise.(**D**) DPA sample decoding accuracy of the APC neurons in the dual-task trials with the No-go distractor.(**E**) Same as (D), but in trials with the Go distractor.(**F**) Correlations between sample decoding of APC activity and dual-task performance.Above: Averaged DPA sample decoding accuracy of the late delay period (left) and correct rate in DPA performance (right) under different conditions of distractors.***, p < 0.001, one-way ANOVA.Below: Correlations of DPA sample decoding accuracy and correct rate in bootstrap resamples.Statistics, the Pearson correlation coefficient.

## Discussion

Our work causally demonstrated that the APC was critical in the active maintenance in olfactory working memory. Using optogenetic manipulation in a series of behavioral tasks that temporally isolated retention of sensory information during the delay period from decision making, we demonstrated that the APC delay activity was critical for active information maintenance in working memory with or without distraction. The control experiments demonstrated that the behavioral defects of optogenetic perturbation during the delay period were due to impaired information maintenance. The APC population activity exhibited mutual information and decoding power for the odor samples during the odor-delivery and delay periods.Therefore, although the APC activity is constantly updated by ongoing sensory inputs, this sensory cortex is resistant to overwriting and can maintain information through population activity in the WM tasks, even with a distracting task during the delay period. This is contrary to the prediction by the theory of passive reflection from higher associative areas and supports the critical role of the APC in active maintenance of information for olfactory WM. The functional role of other sensory regions (Ghosh et al., 2011; Miyamichi et al., 2011; Sosulski et al., 2011) and the potential changes in interaction between the APC and the mPFC (Liu et al., 2014) during learning of the WM tasks remains to be determined. Our results therefore demonstrated that the APC is a critical node in the cortical network responsible for information maintenance in the WM tasks.

### Memory vs.sensory perception

Previous studies debated whether the delay-period activity of sensory regions is actively maintaining information or passively reflecting top-down inputs from higher cortices (Pasternak and Greenlee, 2005) (Colombo et al., 1990; Guo et al., 2014; Harris et al., 2002; Mendoza-Halliday et al., 2014; Seidemann et al., 1998). Until recently we lacked the necessary techniques with millisecond-precision in perturbation of neuronal activity and unambiguous task design to determine: (1) whether the perturbation during the delay-period activity in sensory regions induce behavioral deficits; (2) whether the sustained memory-selective delay-period activity in the sensory areas is robust to intervening distractors.

In this study, we found that optogenetic suppression of the APC activity during the delay period impaired performance in a series of olfactory working memory tasks.Furthermore, electrophysiological recordings revealed that the delay-period activity can code for odor information even with an overlapping distracting task.Thus, the APC is critical for active information maintenance in olfactory working memory.

### Distributed network Interaction

The olfactory pathway downstream from the olfactory bulb is organized in a highly parallel manner, as mitral/tufted neurons in the olfactory bulb project to multiple brain regions, including the APC, the anterior olfactory nucleus, the cortical amygdala, the olfactory tubercle, and the lateral entorhinal cortex (Davison and Ehlers, 2011; Ghosh et al., 2011; Miyamichi et al., 2011; Price and Powell, 1970; Sosulski et al., 2011).Therefore, it was surprising that optogenetic perturbation of just one out of five brain regions can impair olfactory WM performance, suggesting the critical importance of the APC for WM among the parallel olfactory pathways. The functional role of other sensory regions (Davison and Ehlers, 2011; Ghosh et al., 2011; Miyamichi et al., 2011; Price and Powell, 1970; Sosulski et al., 2011) and potential interaction between the APC and the mPFC (Liu et al., 2014) in learning and well-trained phases of WM tasks remain to be determined.

### Dynamic transfer of the functional role from mPFC to APC through learning

Previously we showed that the delay-period activity of the medial prefrontal cortex (mPFC) was only important during the learning but not the well-trained phase in the olfactory WM task (Liu et al., 2014). Therefore it is pertinent to examine the roles of different brain regions in learning and well-trained mice. The results of our optogenetic and recording experiments causally demonstrated that the APC delay activity was important in the well-trained phases.Thus, the functional role of memory maintenance is transferred from the mPFC to the APC through learning. The underlying mechanisms of the transfer and the potential involvement of other brain regions remain to be determined.

The APC neuronal activity is associated with olfactory perception, but it also varies depending on brain states, task design, or learning experience (Courtiol and Wilson, 2016). For this reason the APC has long been suggested to be an associative sensory region (Bekkers and Suzuki, 2013; Courtiol and Wilson, 2016). Consistent with this notion, our results causally demonstrated the importance of the APC activity beyond sensory perception. In summary, our results underscore the importance of olfactory sensory cortex in memory maintenance and behavioral choices beyond immediate sensory processing.

## Acknowledgments

We thank Dr.M.P.Stryker, B.Richmond, R. Desimone for communication on head-fixed mice preparation and behavioral design; Dr.G.P.Feng for VGAT-ChR2 mice; Dr.G.Laurent for communication on population analysis; Dr.L.P.Wang for help in optogenetics; Dr.L.N.Lin for help in tetrode drive manufacturing; Dr.Y.Dan and J.F.Erlich for communication on spike sorting; Drs.Z.L.Qiu, M.M.Luo, H.L.Hu and X.Yu for help on transgenic mice; Dr.A.K.Guo for help in PID experiments; Imaging facility of ION and Dr.Q.Hu for help in imaging experiments; Dr.Y.F.Li for drawing the summarizing schematic; and Dr.M.M.Poo for critical comments on the manuscript. The work was supported by the National Science Foundation for Distinguished Young Scholars of China (31525010, to C.T.L.), the General Program of the Chinese National Science Foundation (31471049), the Instrument Developing Project of the Chinese Academy of Sciences (Grant No.YZ201540), the Key Research Program of Frontier Sciences of the Chinese Academy Sciences (QYZDB-SSW-SMC009), the Key Project of the Shanghai Science and Technology Commission (No.15JC1400102, 16JC1400101), the China–Netherlands CAS-NWO Programme: The Future of Brain and Cognition (153D31KYSB20160106).

## Author Contributions

X.Z., W.Y and C.T.L.conceived and designed the experiments.A.C.and S.D.gave critical suggestions for the experiments.X.Z., W.Y and Y.C.performed behavioral, electrophysiology and optogenetic experiments.H.F.constructed the recording electrodes and managed transgenic mice.R.H.and Z.C.performed mice surgery and helped in blind design.X.Z., W.Y, and C.T.L.analyzed data.W.W., A.C., and S.D.helped in data analysis.X.Z.and C.T.L.wrote the paper.

## STAR Methods

### CONTACT FOR REAGENT AND RESOURCE SHARING

Further information and requests for resources and reagents should be directed to and will be fulfilled by the Lead Contact, Chengyu T.Li.

### EXPERIMENTAL MODEL AND SUBJECT DETAILS

#### Animals

All experiments were performed in compliance with the animal care standards set by the U.S.National Institutes of Health and have been approved by the Institutional Animal Care and Use Committee of the Institute of Neuroscience, Chinese Academy of Sciences (Shanghai, China).B6.Cg-Tg(Slc32a1-COP4*H134R/EYFP)8Gfng/J mice, commonly referred to as VGAT-ChR2 mice were used in optogenetic experiments, and littermates of the same sex were used as control. All mice were healthy male, group housed, of age 8-12 weeks and 20-30g in weight at the start of training. For all experiments, individual animal are presented as individual data points, otherwise the sample size (n) are shown in the corresponding figures. We ensured each group in behavior studies included at least 10 mice, which had been shown to be suitable for detecting effects of optogenetic manipulations in comparable working memory tasks(Liu et al., 2014).

### METHOD DETAILS

#### Behavioral setups

We utilized the DNMS task, DPA task and dual-task as described previously (Liu et al., 2014; Han et al., 2018) or described as follows. In brief, the olfactometry apparatuses was enclosed in sound-proof training boxes. An embedded system customarily built around a PIC Digital Signal Controller (dsPIC30F6010A, Microchip) was used to control the olfactory cue and water delivery by switching solenoid valves and to detect lick responses with an infra-red beam break detector. The air-and-odorant-mixture nozzle/lick-port/beam-breaker assembly were 3D-printed and placed in front of the mouse. The air flow rate is controlled at 1.35L/min during and between odor deliveries.N-butanol (1:500; B7906, Sigma-Aldrich), propyl formate (1:500; 245852, Sigma-Aldrich), ethyl acetate (1:500; 270989, Sigma-Aldrich), 3-methyl-2-buten-1-ol (1:500, W364703, Sigma-Aldrich), 1-pentanol (1:500, 398268, Sigma-Aldrich) and butyl formate (1:500; 261521, Sigma-Aldrich) were used at specified concentration in mineral oil (v/v; O1224, Fisher Scientific). For the multiple-sample DPA task, the stock odorants are exposed to individually controlled air flow, and the odor-mixtures were mixed to the air at 1:10 concentration (v/v). The concentration of odors were measured by a photoionization detector (200B miniPID, Aurora Scientific Inc.), and the concentration during delay duration reduced to that of the baseline level within 1s after valve shut-off. Behavior events and timings were simultaneously sent to and recorded by a computer using customarily written software.

#### Behavior training

The DNMS task was carried out as described previously(Liu et al., 2014).Briefly, a sample olfactory stimulus was presented for 1 second at the start of a trial, followed by a delay duration of 4-40 sec (mice need to retain the information of the first stimulus (sample) during the delay duration), then a test olfactory stimulus for 1 second, identical to (in match trials) or different from (in non-matched trials) the sample. After a 1-second pre-response-delay, mice were trained to lick in the 0.5s response window only in non-matched trials.*Hit* or *False alarm* was defined as detection of lick events in the response window in a non-match or match trial, respectively.Similarly, *Miss* or *Correct rejection* were defined as absence of a lick event in the response window in a non-match or match trial.A reward of 5μl water was triggered immediately only after *Hit;* mice were neither rewarded nor punished following other responses. Mice were allowed to perform up to 300 trials each day, a consecutive combination of 10 miss and correct rejection trials also triggers the end of the session; only the trials within well-trained performance windows (no less than 80% correct rate within consecutive 40 unperturbed trials) were included for data analysis, unless stated otherwise. In the increasing delay duration experiment, the mice were trained to perform the DNMS task with 5s delay to the well-trained criterion, then the delay duration was increased every day; the first 100 trials was included for the analysis to assist parallel comparison. An inter-trial interval twice as long as the delay duration separated consecutive trials.

In the DPA task, one of two sample odor was presented for 1 second at the start of a trial, followed by a delay duration of 13 sec, then one of two different test odors for 1 second. One of the sample odor and one of the test odor formed a rewarded pair, while the other two odors formed another rewarded pair. After a 1-second pre-response-delay, mice were trained to lick in the 0.5s response window only in paired trials.*Hit* or *False alarm* was defined as detection of lick events in the response window in a paired or non-paired trial, respectively.Similarly, *Miss* or *Correct rejection* were defined as absence of a lick event in the response window in a paired or non-paired trial.A reward of 5μl water was triggered immediately only after *Hit;* mice were neither rewarded nor punished following other responses. Mice were allowed to perform up to 300 trials each day, a consecutive combination of 10 miss and correct rejection trials also triggers the end of the session; only the trials within well-trained performance windows (no less than 80% correct rate within consecutive 40 unperturbed trials) were included for data analysis, unless stated otherwise. An inter-trial interval as long as the delay durations separated consecutive trials.

The multiple sample DPA (MS-DPA) task is like the DPA task described previously, except that the number of candidate sample odors were increased to 6; three of the samples and one test formed rewarded pairs (e.g., S1, S2, S3 and T1), and the other 3 samples and the other test formed remaining rewarded pairs (e.g., S4, S5, S6 and T2). The delay duration in the MS-DPA task is 5 seconds.

In the dual-task, a secondary Go/No-go task was inserted into the delay duration of the DPA task. The olfactory cue delivery of the Go/No-go task started at the third second into the DPA task delay period and was presented for 0.5 second. After a 0.5-second pre-response-delay, mice were trained to lick in the 0.5s response window after the Go stimulus.*Hit* or *False alarm* of the Go/No-go task was defined as detection of lick events in the response window in a Go or No-go trial, respectively.Similarly, *Miss* or *Correct rejection* were defined as absence of a lick event in the response window in a Go or No-go trial.A reward of 5μl water was triggered immediately only after *Hit;* mice were neither rewarded nor punished following other responses. The sample and test stimuli of the DPA task and the stimulus of the Go/No-go task are arranged independently in a pseudo-random fashion. Mice were allowed to perform up to 288 dual-task trials each day, a consecutive combination of 10 miss and correct rejection trials also triggers the end of the session; for the well-trained phase studies, only the trials within well-trained performance windows (no less than 80% DPA correct rate within consecutive 40 unperturbed trials) were used for data analysis; for the learning phase, all trials are included in the analysis, unless stated otherwise. An inter-trial interval as long as the delay duration of the DPA task separated consecutive trials. The Go/No-go task is omitted in one third of all trials. The performance in the 4th day of the dual-task learning stage was used to estimate the dual-task interference (**Figure 4E**).

Mice were water restricted for 48 hours before start of training, followed by habituation, shaping and learning phases, before well-trained optogenetic sessions. In habituation phase, mice were head-fixed in the olfactometry apparatus for 2hrs and allowed to lick water from the water port, encouraged by program controlled water delivery. Typically in 1 to 2 days, mice could learn to spontaneously trigger more than 100 water rewards. In the shaping phase, only non-match or paired trials were presented, and water will be pseudo-randomly and programmatically delivered after 1/3 of the *Miss* responses to encourage task engagement. Shaping phase ended once the mice could perform more than 32 *Hit* responses in consecutive 40 trials (80% in performance). In learning phase, both match and non-match, paired and non-paired trials were presented, and each four consecutive trials that consist of 4 balanced sample-test-reward combinations were shuffled. Learning phase ended when mice could perform more than 32 *Hit* / *Correct Reject* responses in consecutive 40 trials (80% in performance). The performance (referred to as “correct rate” in labels of figures) was defined as the total fraction of hit and correct rejection response. The sensitivity index (d’) was defined as *d*’ = *norminv(Hit rate)-norminv (False choice rate)*. The lick efficiency was defined as the number of rewarded lick divided by the sum of rewarded and unrewarded lick in the response window.(Liu et al., 2014).

For all experiments with blind design, R.Q.Hou, H.M.Fan and Z.Q.Chen labeled the mice with unique numbers without revealing the genotype; they would not participate in behavior or optogenetic experiments. The genotype of mice would only be revealed after the experiments and statistical analysis of individual mice had been finished. Based on the blind design and the task design in which all mice participate the same behavior studies in identical sequences, there is no need for further randomization of samples in this study.

#### Stereotaxic implantation of optical fiber

Implantation of optical fiber were similar to previous description(Liu et al., 2014).Briefly, Mice were anaesthetized and placed in a stereotaxic instrument (Stoelting Co.). After removal of the scalp, periosteum and other associated soft tissues, a custom designed steel plate was fixed on skull by tissue adhesive (1469SB, 3M) and dental cement. Craniotomies of roughly 1 mm in diameter were made bilaterally above the anterior piriform cortex (APC). For optical fiber implantation, two optic fibers (200 μm in diameter, Leizhao Biotech., China) with ceramic ferrule were implanted at A.P.1.78 mm, M.L.2.60 mm and D.V.3.40 mm.A thin layer of silicone elastomer (Kwik-Sil, WPI) was applied to protect the brain tissue, then dental cement were applied to connect the skull, plate and optical fibers for structural support. Antibiotic drug (ampicillin sodium) was i.p.injected for 3 consecutive days after surgery.

#### Optogenetic experiments

Optogenetic experiments were performed as described previously(Liu et al., 2014).Briefly, an optical patch cable (200 μm in diameter, N.A.0.37) was used to connect the implanted optic fiber (through a ceramic sleeve) to a laser source (BL473T3-50FC, SLOC, China). Laser power was programmatically controlled by the PIC microcontroller through analog voltage input. Output laser power was measured with a laser power meter (LP1, Sanwa, Japan) and compensated for optical power loss in fiber implant and coupling attenuation (determined before surgery). In optogenetic experiments with laser power ramp, the dynamics were further calibrated by an oscilloscope.

In the DNMS task optogenetic sessions (**Figures 1H-1N, 2B-2D**, and **S1**), the optogenetic perturbation was arranged in an interleaved one-trial-on, one-trial-off fashion; each session started with a laser-off trial. In the DPA task and dual-task optogenetic sessions (**Figures 4B, 4F and S2**), the optogenetic perturbation was arranged in an interleaved one-block-on, one-block-off fashion, and each block consisted of 24 trials; each session started with a laser-off block.

#### Immunostaining and imaging

After the conclusion of experiments, mice were deeply anesthetized with sodium pentobarbital (120 mg / kg) and then perfused transcardially with 20 mL 0.9% NaCl solution followed by 20 mL paraformaldehyde PBS solution (PFA, 4%, w/v). The brains were fixed in PFA solution overnight and sectioned. Slices were washed with PBS, fluorescence-enhanced with the anti-GFP antibody (FITC, ab6662, Abcam), incubated with DAPI (C1002, 1:1000, Beyotime), and were mounted with DAPI. Fluorescence images were then obtained with an epifluorescence microscope (Eclipse 80i, Nikon) with 10X (CFI Plan Apo Lambda, 0.45 N.A.) or 20X (CFI Plan Apo Lambda, 0.75 N.A.) objective lens, and analyzed with ImageJ software (NIH, U.S.).

#### Surgical implantation of (op-)tetrode microdrive

The implantation procedure was similar to previous report(Liu et al., 2014).Briefly, all surgical tools and the microdrive were sterilized by ultraviolet radiation for more than 20 minutes before implantation. One cranial window of 0.5 by 1 mm was made in either hemisphere centered at A.P.+1.42mm, M.L.2.5mm, and electrodes were lowered at D.V.3mm to target the APC. Tissue gel (1469SB, 3M) and dental cement were carefully applied to cover the exposed brain tissue and to fix the microdrive. Antibiotics (ampicillin sodium, 20 mg/mL, 160 mg/kg b.w.) was injected for three consecutive days after surgery. Behavior training started 7 to 14 days after that. Recording of behaving mice were made with electrodes lowered for approximately 50 μm each day after the last day of shaping.

#### Electrophysiology recording

Wide band signals (0.5-8000 Hz) from all tetrodes were amplified (× 20000) and digitized at 40 kHz with the Multichannel Acquisition Processor (Plexon Inc.) and all data were saved to hard-disks. Spike events detection and sorting was performed offline as described in previous paper(Liu et al., 2014).Briefly, offline spike detection was performed with Offline Sorter (Plexon Inc.). Raw signals were filtered in 250~8000 Hz to remove field potentials. Signals larger than five times of standard deviation recorded on any recording site of tetrode were considered as spike events. Principle component analysis (PCA) was performed for tetrode-waveforms to extract the first three principle components (PCs) explaining the largest variance.Then, contour or valley-seek clustering provided by Offline Sorter was performed in 2D or 3D feature space (including principle components) of waveforms. Single units was included only if there were no more than 0.15% of spikes occurred within 2 ms refractory period and the averaged firing rate was higher than 2 Hz. All recording sites were further confirmed by passing current (50μA, 100ms, 1Hz x 5 pulses) through the electrodes, and verified with DAPI staining and immunochemistry 1 day after the lesion.

### QUANTIFICATION AND STATISTICAL ANALYSIS

All further analyses were performed by custom-written codes with MATLAB (Mathworks) and Java (Oracle). Statistical significance was defined as p < 0.05 unless noted otherwise.

#### Optogenetic perturbation

The genotype of mice would only be revealed after the experiments and statistical analysis of individual mice had been finished. Then the behavior performance in trials without and with optogenetic perturbation were calculated separately for experiment group and control group. For the DNMS task, one-way ANOVA (MATLAB function *anovan())* of correct rate change (Δ correct rate) was performed for ChR2 mice across all delay durations.Δ values were defined as the values of laser-on trials minus that of laser-off trials. Wilcoxon rank-sum test (MATLAB function *ranksum())* were performed for Δ values between control and ChR2 mice in all the optogenetic tasks. Then paired Student’s t-test (MATLAB function *ttest())* were performed for ChR2 mice between laser-on and laser-off trials.

#### General linear model

The DNMS task trials can be defined by a combination of task parameter values, including sample odor, test odor, match or non-match relationship, mice genotype, laser-on or laser-off per trial, delay duration (transformed to an exponential time-decay value based on **Figure 1D**), and the perturbation design of no perturbation, perturb baseline or perturb delay.A subset of aforementioned task parameters, expressed as vector *X*, and a coefficient vector *b*, defined a linear combination *XTb*. The trials corresponding to the combination of X were group together, and the averaged correct rate of the trials were collected as response value μ. The model is *μ= XTb*.A series of 7 models were systematically generated as descripted in **Figure 3A**, and fitted with MATLAB function *fitglm()*. The resulting model and corresponding r-squared and Akaike information criterion (AIC) value were examined and the one (#6) with the lowest AIC value were chosen for further analysis. The combination of each *X* and *b* in the model provided the numerical relationship between a task parameter and the averaged change in correct rate in **Figure 3D.**

#### Spike count and activity heat map

Averaged firing rates of individual neurons during different olfactory sample were binned in 200ms (**Figures 6B** and **S3**). The baseline period was defined as the one second before onset of the sample odor delivery. Firing rates from baseline of each trial were averaged to form the baseline activity vectors for each neuron. Mean and standard deviation of this baseline activity vector were used to convert averaged firing rates into Z-scores. Activity of all neurons following different sample odors were sorted by the differences between sample odors during the delay period and plotted as heat map using the “jet” color-map defined in MATLAB.

#### Mutual information

Briefly, the spike counts for each neuron during task trials were binned in 500ms windows moving at 100ms resolution, and transformed to firing rates, then grouped according to the sample odor. The distribution of firing rates within a time window for the same sample odor were fitted to a Gaussian distribution with MATLAB *fitdist()* function. The probability of a response rate r, *P[r]*, was defined by the probability density function of the fitted Gaussian distribution. The probability of a stimulus *P[s]* was defined as the fraction of trials with sample s in all the trials. The conditional probability *P[r|s]* represented the probability of a response rate *r* given that sample was *s*. The mutual information for the sample of a neuron in a task was thus calculated as

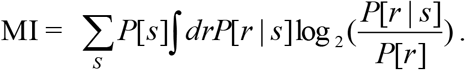

#### SVM decoding

The population decoding analysis is performed with LIBSVM library (https://www.csie.ntu.edu.tw/~cjlin/libsvm) according to the documents.Briefly, the spike counts for each neuron during task trials were binned in 500ms windows, and grouped by sample odors. Only neurons with more than 15 well-trained trials for each sample odor were included to minimize overfitting. The firing rates for all neurons were normalized to the [-1, 1] range. For each repeat, firing rates of all neurons in 30 bootstrap trials for each sample odor were selected to train the support vector machine (SVM) with the *radial basis function* (RBF) kernel, and one unselected trial for each sample odor were used to test the classification accuracy. Averaged decoding accuracy was obtained with 500 repeats.A good combination of the two parameters for the RBF kernel, *c* and γ, were grid-searched using exponentially growing sequences in range of 2^[-5, 5] and 2^[-10, 0], respectively. For the shuffled control, the procedure was repeated 1000 times with the nominal sample odor for each trial randomly redistributed. For the study of correlation of behavioral performance and population neuronal decoding (**Fig.7F**), the decoding accuracy was obtained using averaged firing rates from 1 sec before the test onset to the test offset, to reflect the neuronal activity critical to decision making for the DPA task.

### DATA AND SOFTWARE AVAILABILITY

#### Data availability

The data that support the findings of this study are available from the corresponding author upon reasonable request.

#### Code availability

The C and Java source code for the dsPIC based mice training system (Han et al., 2018) can be obtained at: https://github.com/wwweagle/. Other computer code that was used to support the findings of this study are available from the corresponding author upon reasonable request.

---

**Figure S1.**
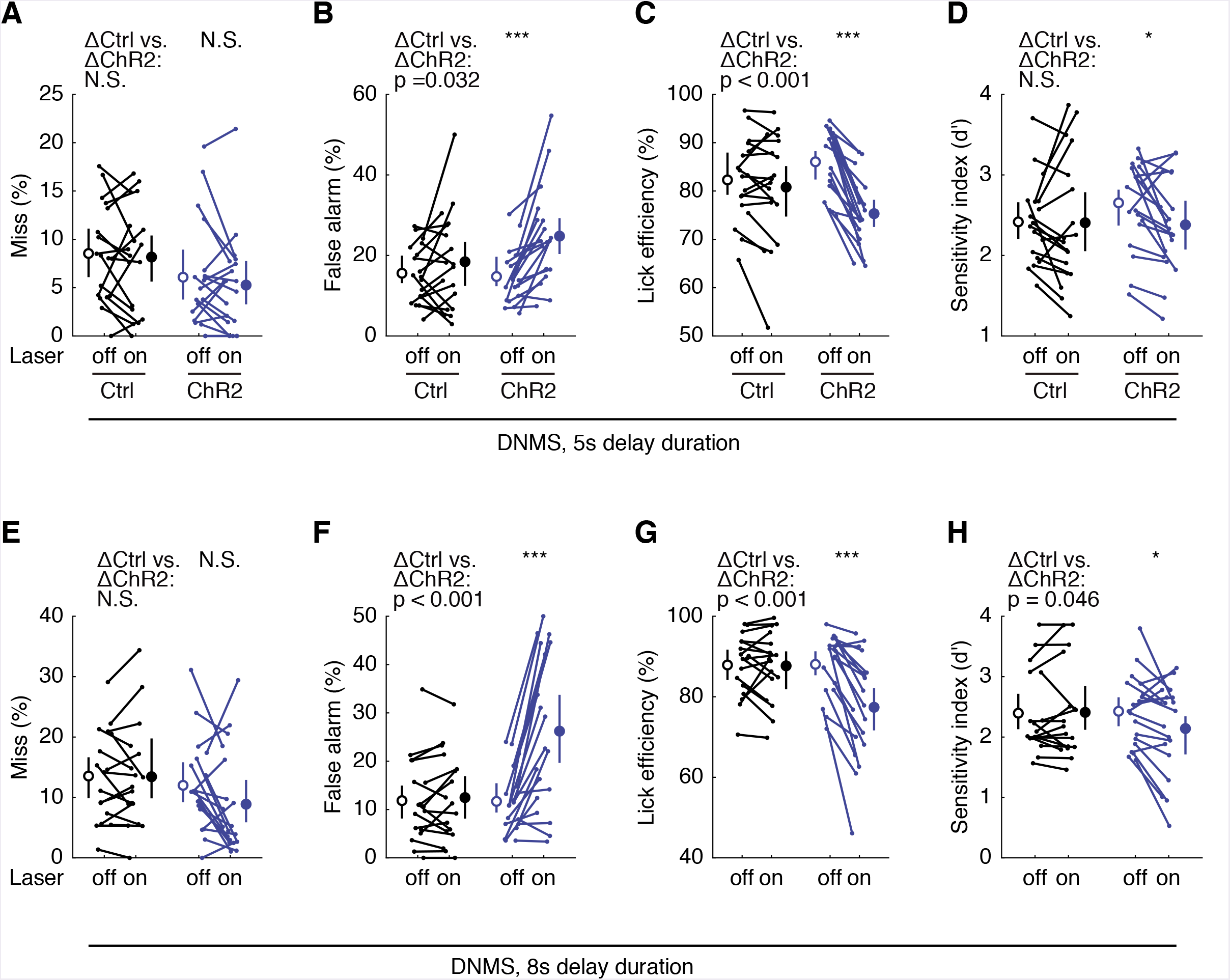
The APC activity is important for the DNMS performance. (**A-D**) Miss rate, false alarm rate, lick efficiency and sensitivity index (d’) following the APC delay-period suppression in DNMS task with 5s delay. Statistic above control groups, the Wilcoxon Rank-Sum test; numbers above ChR2 groups, Student’s paired *t*-test; ***, p < 0.001, *, p = 0.010, error bars: 95% confidence interval of the mean.(**E-H**) Similar to A-D, but in DNMS task with 8s delay.

**Figure S2.**
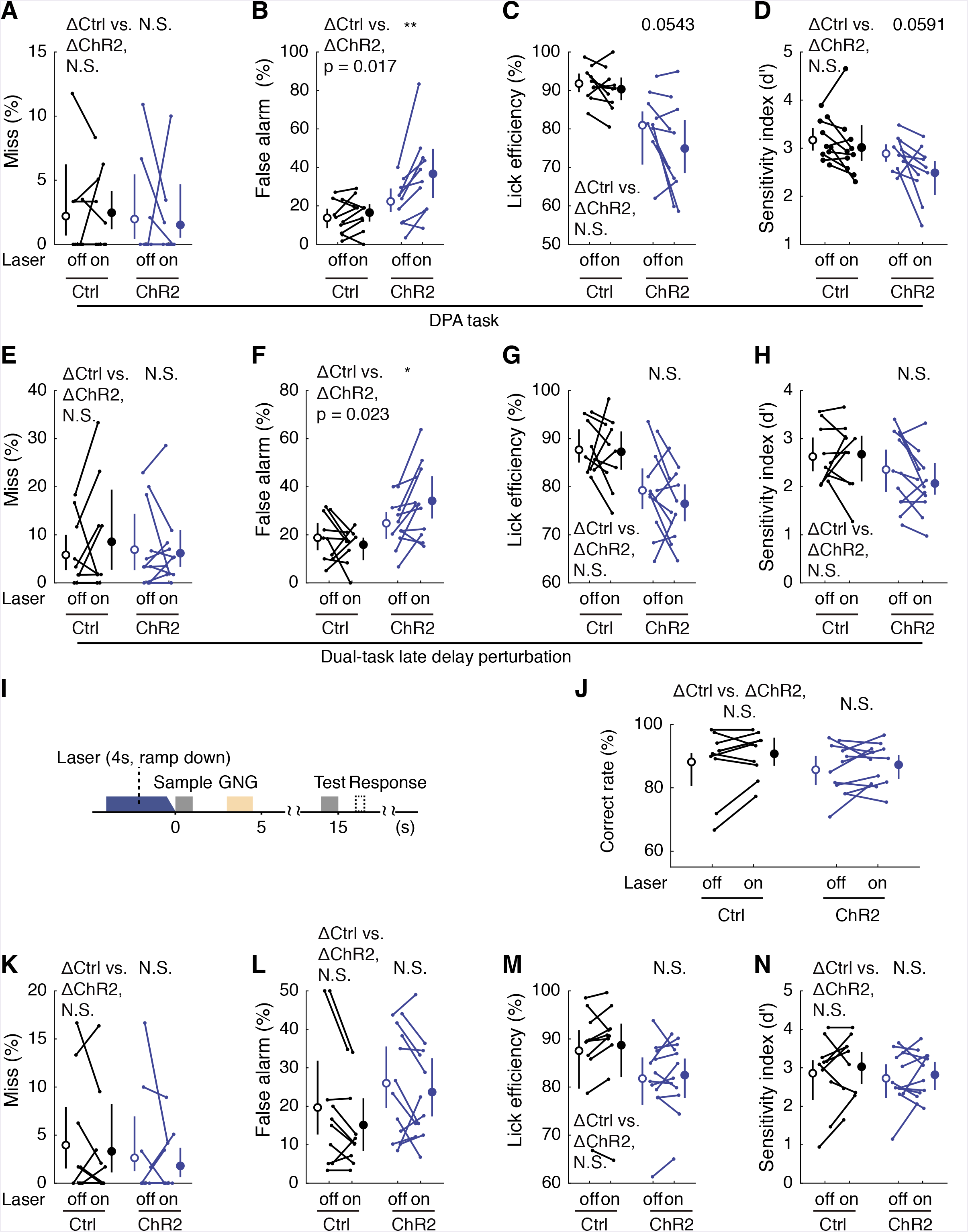
Active memory maintenance by the APC delay activity. (**A-D**) Miss rate, false alarm rate, lick efficiency and sensitivity index (d’) following the APC delay-period suppression in the DPA task. Statistic above control groups, the Wilcoxon Rank-Sum test; numbers above ChR2 groups, Student’s paired *t*-test; **, p = 0.007, error bars: 95% confidence interval of the mean.(**E-H**) Similar to A-D, but in the dual-task.*, p = 0.010.(**I**) Design of the dual-task baseline optogenetic control task.(**J**) Correct rates in the dual-task DPA task following the APC optogenetic suppression during baseline period. Statistic above control groups, the Wilcoxon Rank-Sum test; numbers above ChR2 groups, Student’s paired t-test.(**K-N**)Similar to A-D, but in the dual-task DPA task following the APC optogenetic suppression during baseline period.

**Figure S3.**
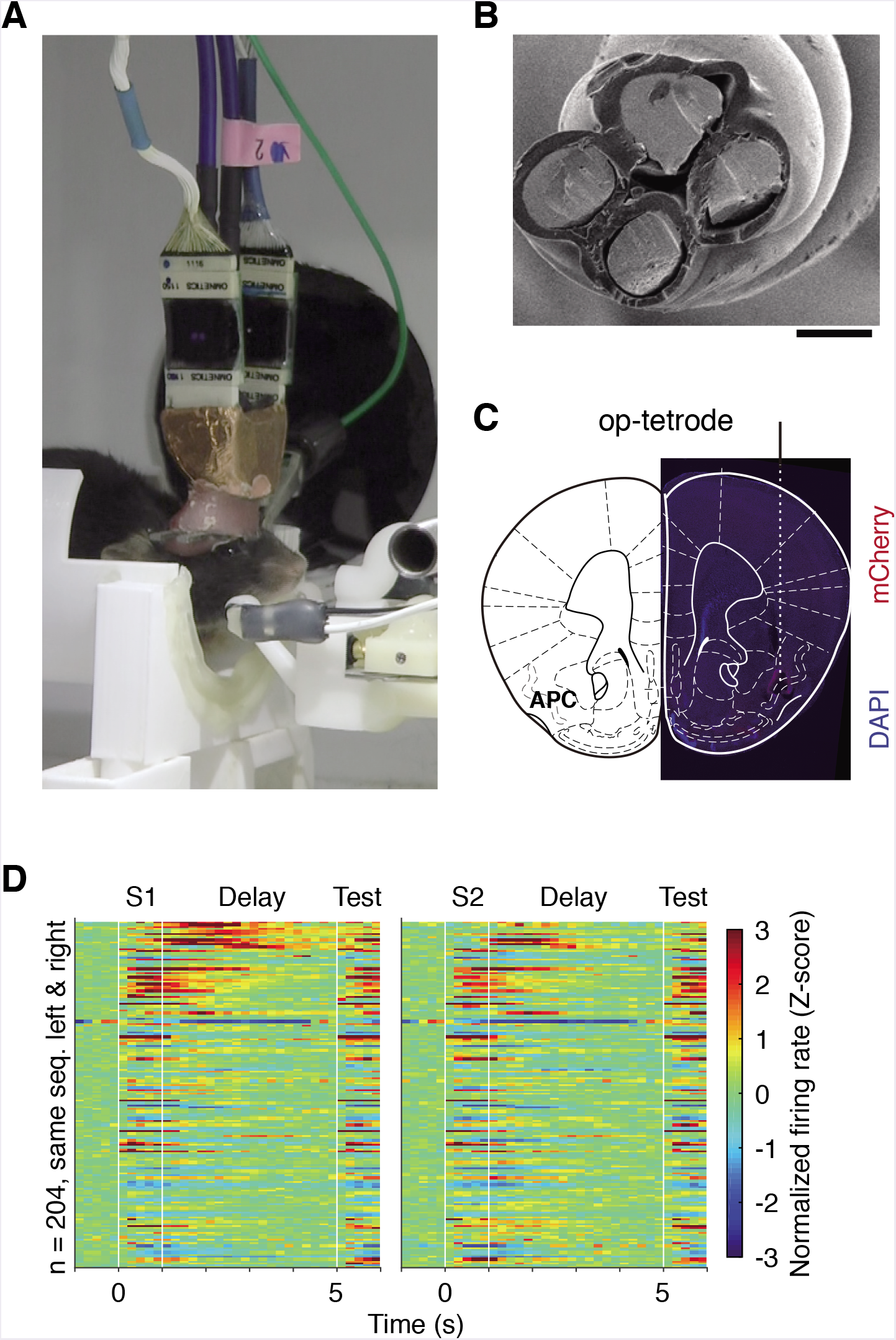
Neural correlates of the APC activity in DNMS task. (**A-C**) Microdrive, tetrode and recording sites. Images of Microdrive (A), tetrode (B, electron microscopy), and recording sites (C, indicated by dashed line).(**D**) Activity of neurons in DNMS task with odor S1 (left) and S2 (right) as sample.Color: firing rates (FR) in Z-scores. Neurons were aligned according to the value of FRS1 – FRS2 during delay period. White gap indicated for the onset of offset of odor delivery

**Table S1.** General linear model coefficients

|  | Estimate | S.E. | T stat | p-value |
| --- | --- | --- | --- | --- |
| Intercept (Constant) | 94.06114 | 0.946678 | 99.35919 | 6.92E-89 |
| Match | -5.60765 | 0.839358 | -6.68088 | 2.43E-09 |
| Delay duration | -0.21587 | 0.066869 | -3.2282 | 0.001778 |
| Laser x Genotype x Memory decay x Perturb delay | -0.35166 | 0.077866 | -4.51622 | 2.04E-05 |

**Table S2.**
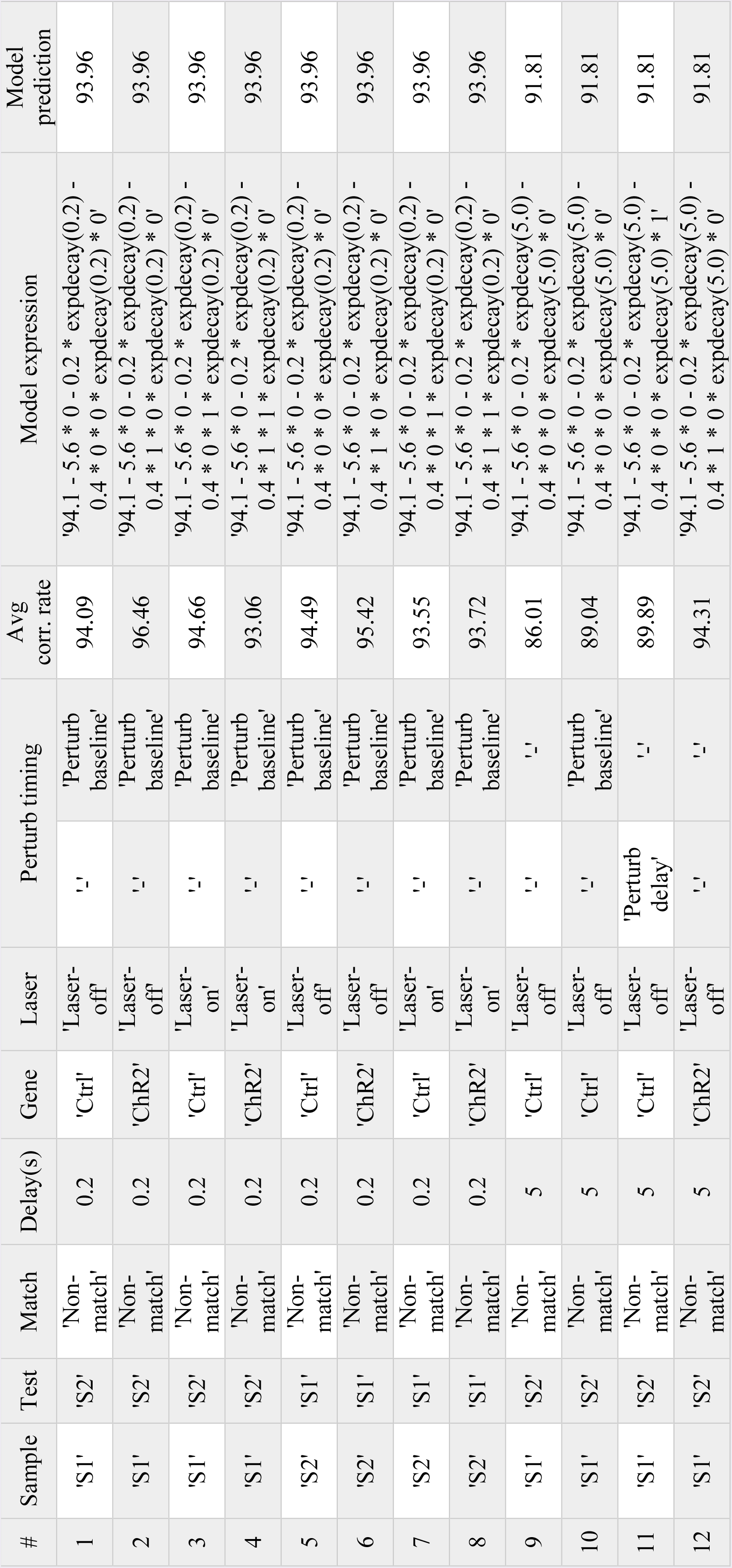
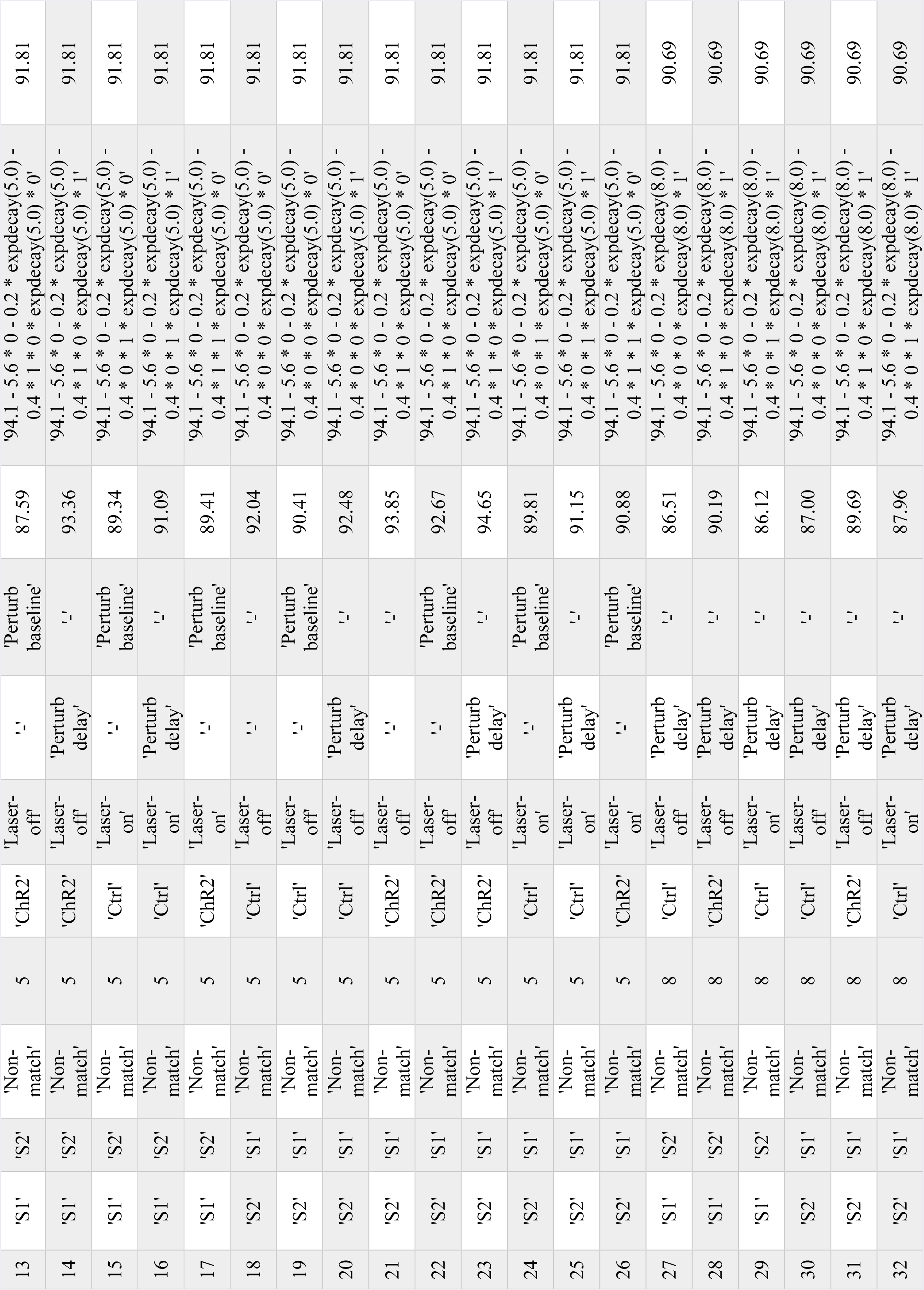
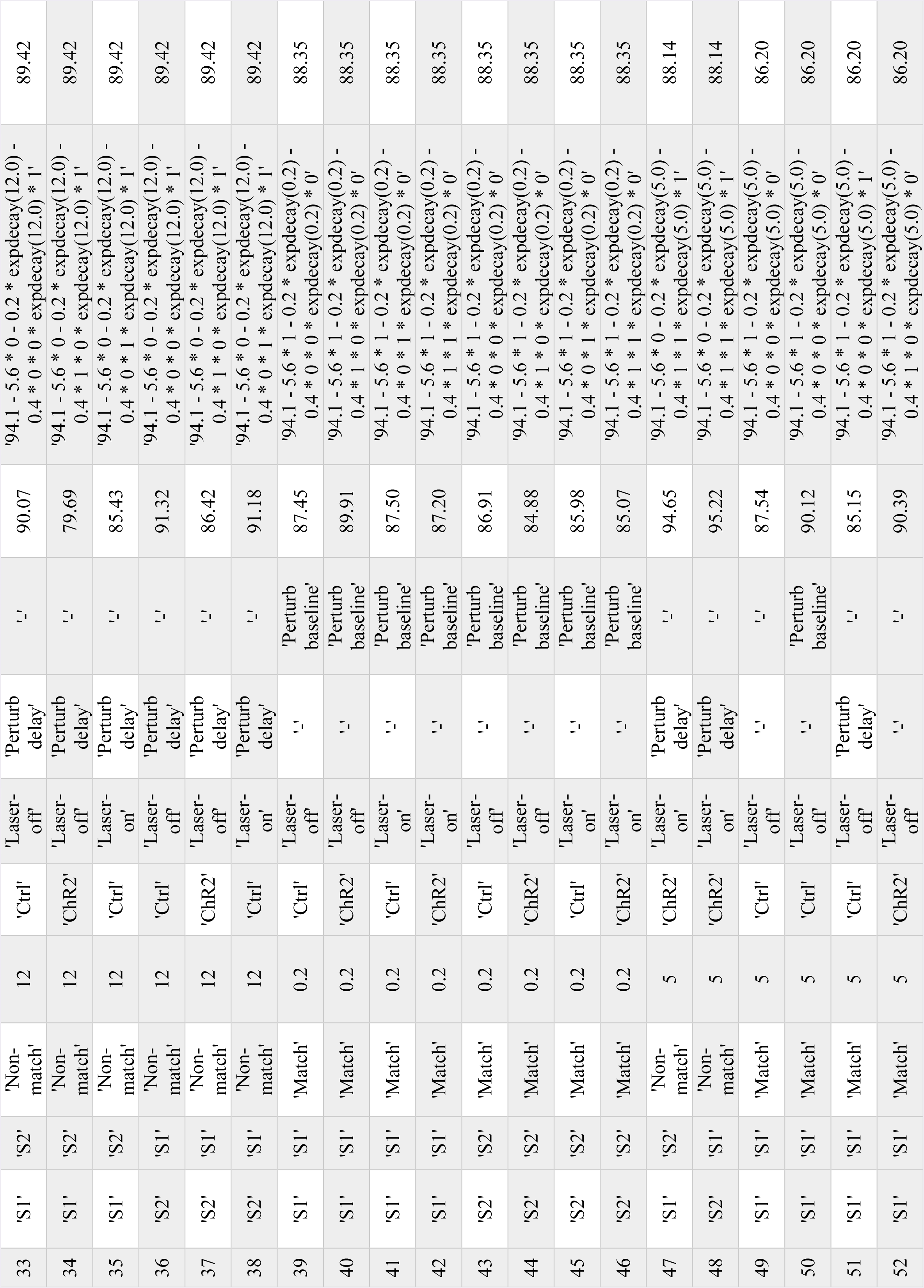
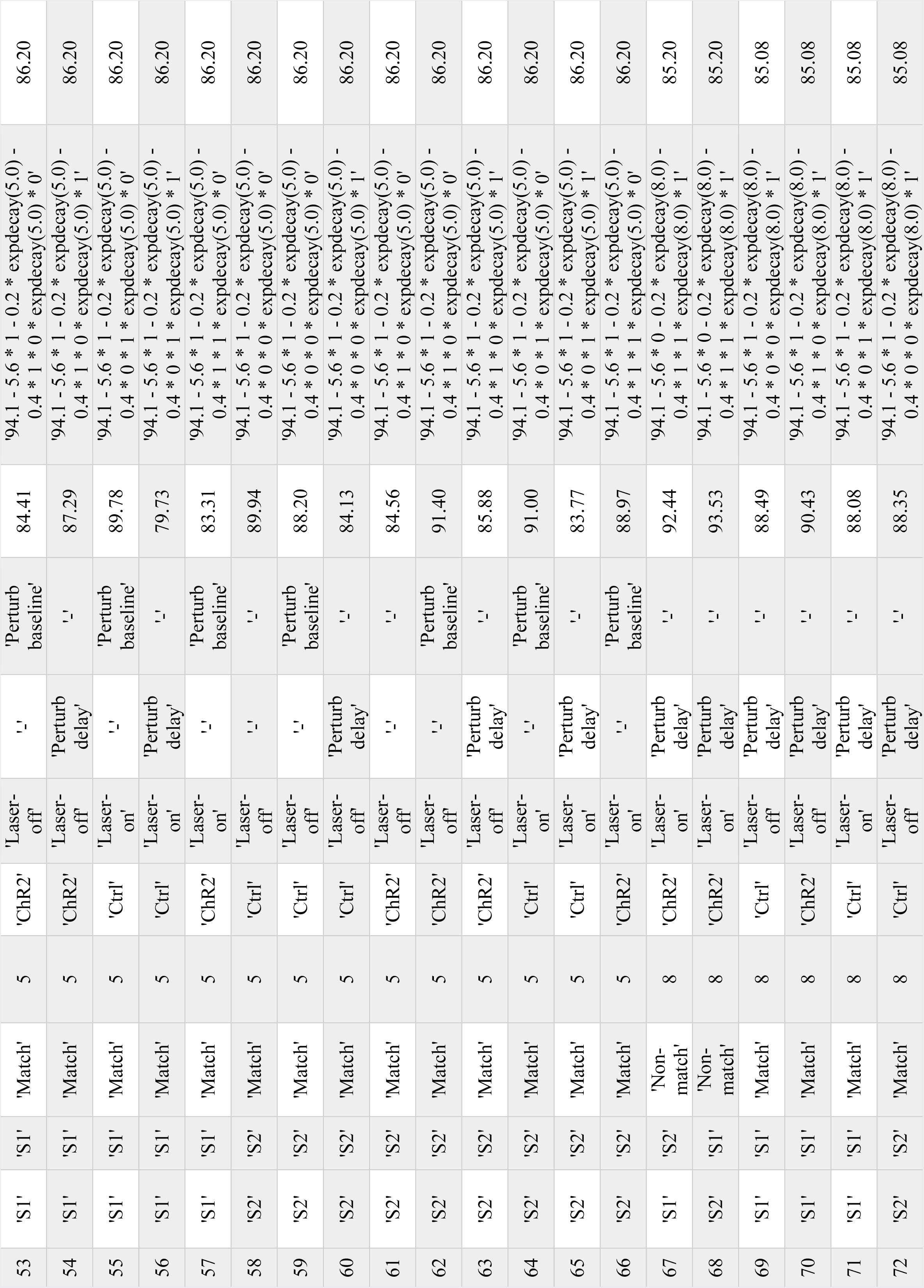
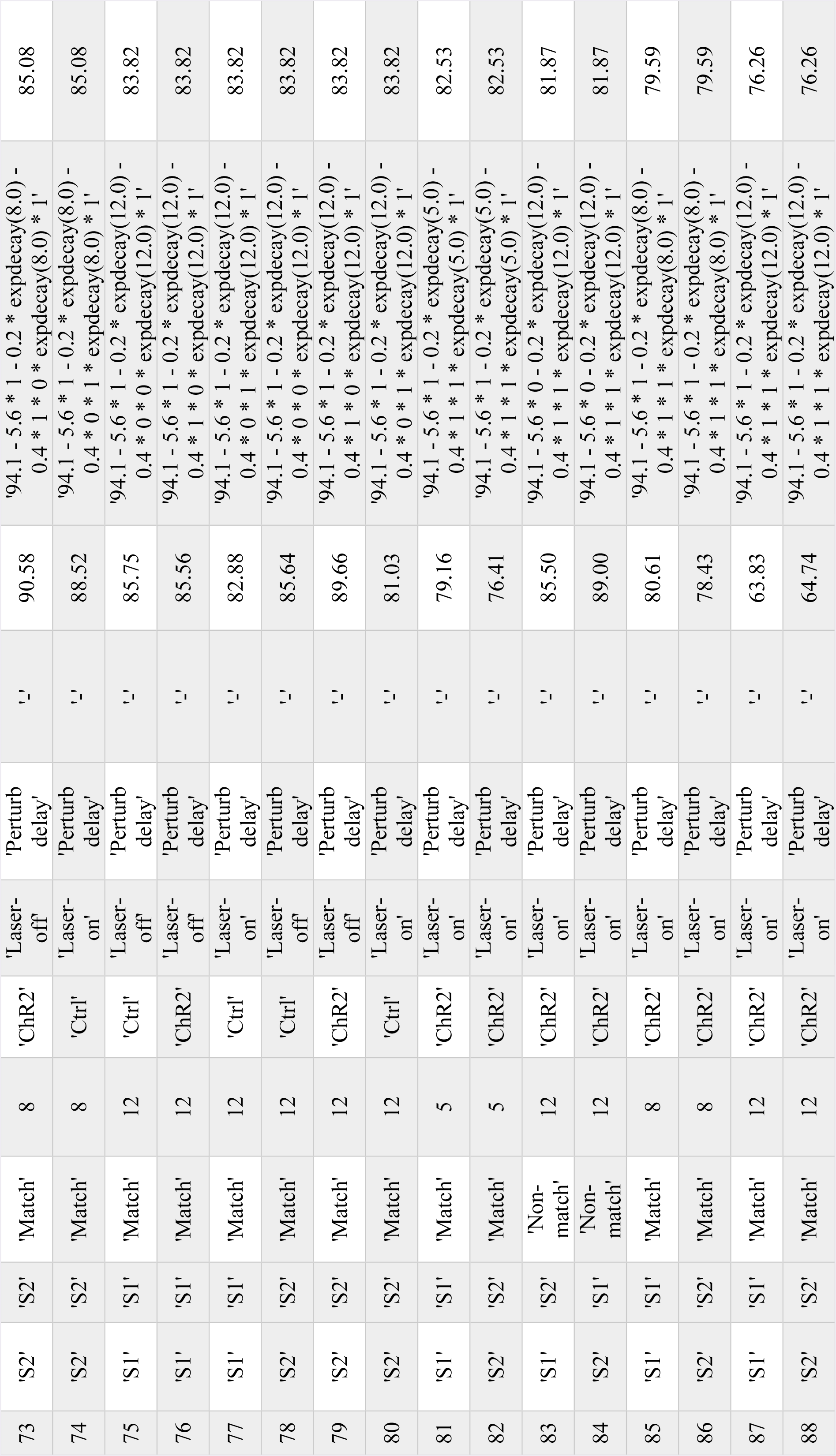
Task parameter combinations and general linear model fit expdecay(t) = 50 - exp(-t / τ) * 50; τ = 21.37

## Supplemental items

